# Oxidative stress leads to Fur-mediated activation of *ftnA* in *Escherichia coli* independently of OxyR/SoxRS regulators

**DOI:** 10.1101/2025.01.22.634231

**Authors:** Valeriia O. Matveeva, Anna D. Grebennikova, Daniil I. Sakharov, Vadim V. Fomin, Ilya V. Manukhov, Sergey V. Bazhenov

## Abstract

Ferritin FtnA is the main scavenger of Fe^2+^ and storage of Fe^3+^ in bacterial cells, together with Dps and Bfr preventing the Fenton reaction and thus protecting the cell from iron-induced oxidative stress. However, until now, it was not known how its expression is regulated under conditions of oxidative stress, and the available evidence was contradictory. To study the regulation of *E. coli ftnA* expression in response to oxidative stress, P_*ftnA*_-*luxCDABE* transcriptional fusion in different strains was used. It has been shown that P_*ftnA*_ is induced after the addition of oxidative stress inducers. The maximum amplitude of this activation did not depend on the presence of functional genes *oxyR* and *soxR* in the cell, but completely disappeared in the absence of *fur*. The response is amplified in the *ftnA* mutant and is diminished in the FtnA-overproducing strain, which indicates that iron sequestration blocks the response. Exposure of a cell to H_2_O_2_ initially inactivates Fur and a number of iron-utilizing proteins, and derepresses iron uptake. This results in an increase in the cellular iron content with the consequent Fur reactivation, which leads to the induction of *ftnA* expression. Thus, oxidative stress leads to P_*ftnA*_ activation, which is mediated by Fur and time-delayed in comparison with OxyR-response.

## 1. Introduction

Ferritin is a spherical protein assembly, which is responsible for oxidizing and storing iron in a cell and is present in a wide range of living organisms (Bai et al. 2015). Typically, ferritin consists of 24 identical subunits, which spontaneously assemble into a full-fledged sphere with a cavity, and free Fe^2+^ is converted to ferrihydrite and stored in the cavity (Honarmand Ebrahimi et al. 2014; Sudarev et al. 2023). In the *Escherichia coli* cells, the protein family of ferritins includes FtnA, FtnB, Bfr, and Dps. These four proteins have significant differences. FtnA is a typical ferritin and acts as a major iron storage (Abdul-Tehrani et al. 1999) and iron depo for re-assembly of iron–sulfur clusters after exposure to hydrogen peroxide or probably another oxidative stressor (Bitoun et al. 2008). FtnB is a distant homologue (33% identity) of FtnA, and they have differences in key amino acids, which casts doubt on its ability to assemble into a spherical complex and to perform ferroxidase activity (Andrews 1998); to date, no one has been able to demonstrate its function experimentally. Bfr contains a heme in its structure and represents an operational storage of iron with the possibility of its recovery through heme-mediated electron transfer (Yao et al. 2012). Finally, Dps forms a sphere of 12 subunits instead of 24 and can bind to DNA and some Fe-utilizing proteins, protecting the key cell components from iron-induced oxidation (Karas et al. 2015).

Proteins from the ferritin family help to protect the cell from iron-induced oxidative stress through the sequestration of free ferrous iron and preventing Fenton reaction. Ferritin FtnA is the major storage of iron in *E. coli* cells, its expression is activated in the presence of Fe^2+^ in a Fur-dependent manner (Massé and Gottesman 2002). Upon binding to Fe^2+^, Fur changes conformation and acquires the ability to bind DNA, most often acting as a transcription repressor. In the work (Nandal et al. 2010), the ability of Fur to act as a transcriptional activator was shown. Fur binds to a distal regulator site upstream of the RNAP-binding site in the promoter of *ftnA* and physically removes the histone-like protein, H-NS, which mediates repression of P_*ftnA*_. The role of Fur as an anti-repressor in the activation of *ftnA* is supported with *in vivo* evidence: (1) *fur* is not required for *ftnA* expression in the absence of *hns*; and (2) *ftnA* expression is not reduced by Fe^2+^-chelation in Δ*hns* (Nandal et al. 2010).

In *E. coli*, the main regulators of the response to oxidative stress are the *oxyRS* and *soxRS* systems (Storz and Imlay 1999). It is known that both systems positively regulate the expression of *fur*, ensuring the activation of its transcription when either hydrogen peroxide or superoxide anion radicals appear in the cell (Zheng et al. 1999). However, the expression of *ftnA* is determined by the amount of Fur-Fe^2+^, not Fur itself (Nandal et al. 2010). So the relationship between the expression levels of *fur* and *ftnA* is less direct than with simple transcriptional regulators like LitR, the amount of which directly affects the expression of target genes (Bazhenov et al. 2021). Works (Blanchard et al. 2007; Fleischhacker and Kiley 2011) suggest that Fur may act as a redox sensor. However, conclusions regarding the regulation of *ftnA* under oxidative stress remain ambiguous: either Fe^2+^ should be oxidized to Fe^3+^ and, consequently, Fur should lose Fe and detach from DNA, reducing *ftnA* expression (Blanchard et al. 2007; Cornelis et al. 2011; Pinochet-Barros and Helmann 2018); or vice versa, Fur must bind to iron released from damaged proteins (Bitoun et al. 2008) and bind to DNA, activating the expression of *ftnA* (Faulkner and Helmann 2011).

In the present work, a series of measurements using whole-cell biosensors was performed to determine the effect of agents that cause oxidative stress on the expression of *ftnA* and *fecA* responsible for the iron storage and uptake in the *E. coli* cells, as well as the roles of the *fur, oxyR*, and *soxR* genes in the regulation of *ftnA* after induction of an oxidative stress.

## 2. Materials and methods

### 2.1. Bacterial strains, plasmids, and growth condition

Bacterial strains and plasmids used in this study are listed in Table 1. The *E. coli* cells were cultivated in liquid or on solid (1.5 % w/v agar is added) lysogeny broth (LB). The medium was supplemented with ampicillin 200 μg/mL, kanamycin 20 μg/mL, streptomycin 20 μg/mL, or their combinations as needed, according to selection markers of used strains.

**Table 1.**
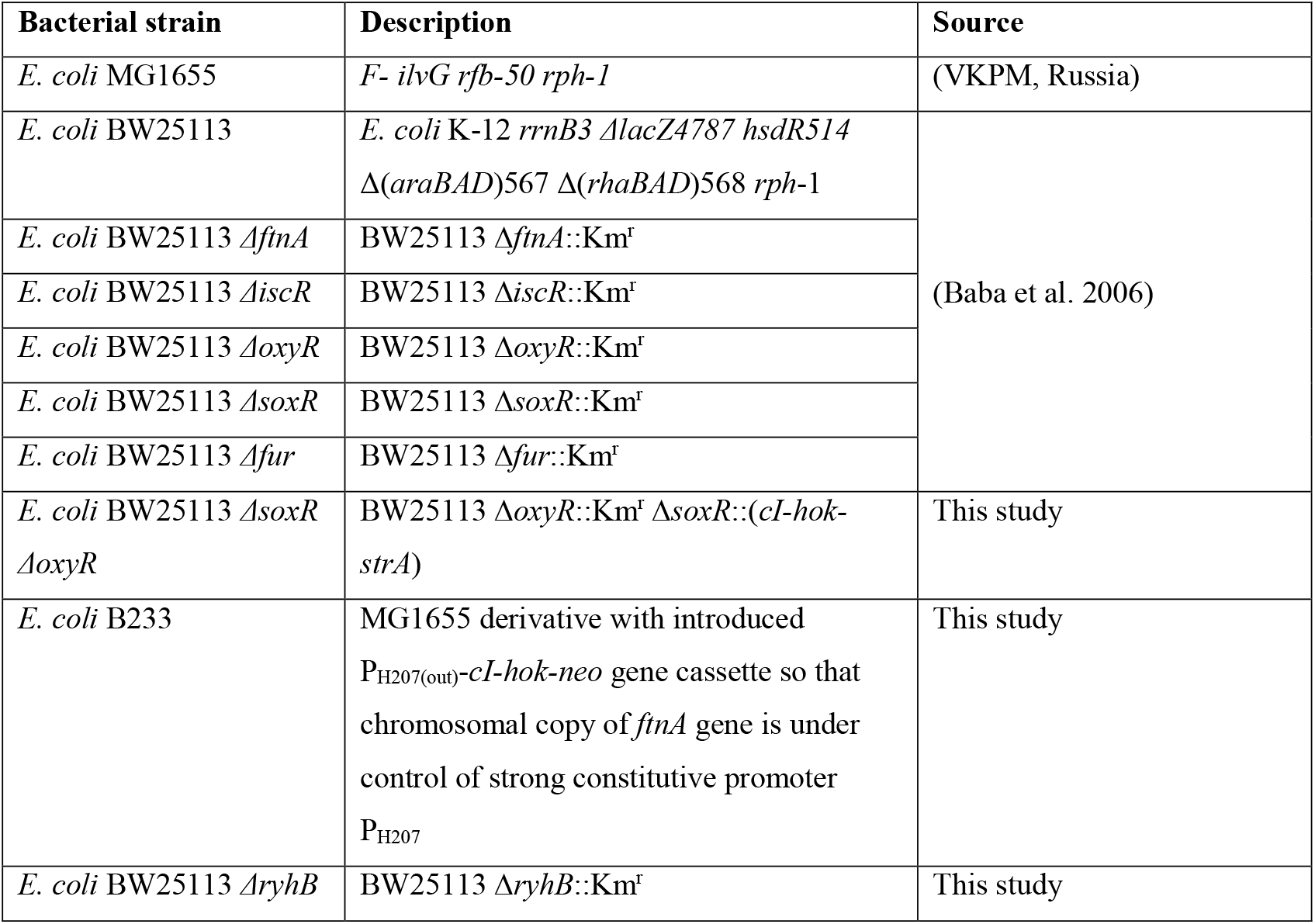
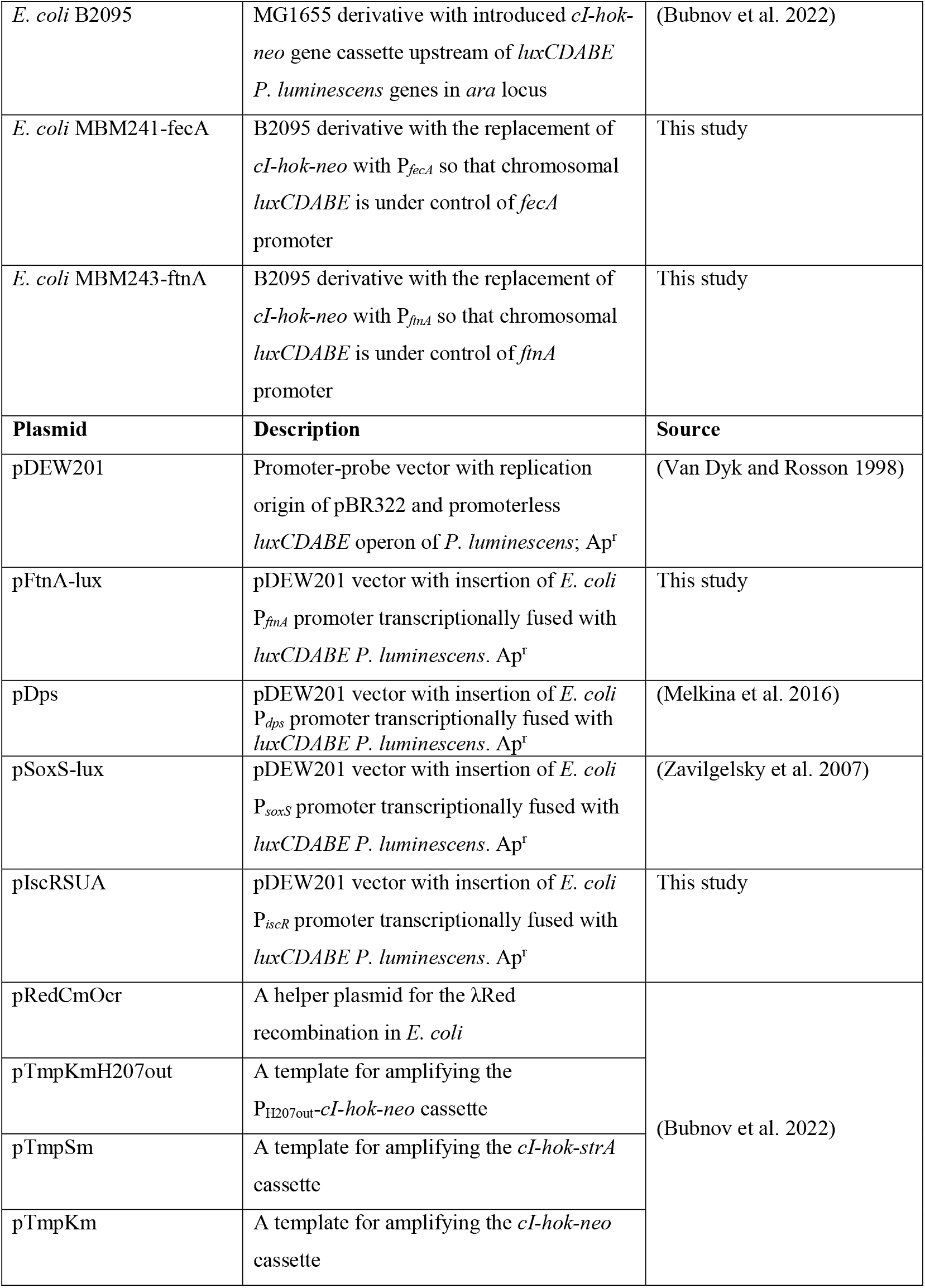
Bacterial strains and plasmids used in this study.

### 2.2. DNA manipulations

Genomic and plasmid DNA extraction, electroelution of DNA from agarose gel, and bacterial transformation were carried out using standard methods (Green and Sambrook 2012). pDEW201 linearization for pFtnA-lux and pIscRSUA plasmids preparation was conducted using EcoRI and BamHI restriction endonucleases. Endonucleases were from SibEnzyme (Russia). Linear fragment had size ∼10 kb, it was purified by electrophoresis and electroelution. To obtain pFtnA-lux biosensor plasmid, P_*ftnA*_ was amplified from *E. coli* MG1655gDNA using primers P1/P2 (Q5 DNA polymerase from NEB, USA, was used) and inserted into the linearized pDEW201 by Gibson Assembly (Gibson et al. 2009). To obtain pIscRSUA biosensor plasmid, P_*iscR*_ was amplified from *E. coli* MG1655 gDNA using primers P3/P4 (Taq polymerase from Evrogen, Russia, was used), subcloned into the pTZ57R/T T-vector using T4 DNA ligase (InsTA cloning kit, ThermoFisher Scientific, USA), and transferred into the pDEW201 vector by EcoRI and BamHI sites. Correct assembly of biosensor plasmids was proved by sequencing with P5 primer. Primers used in the study are presented in Table S1.

### 2.3. Construction of E. coli mutant strains

Chromosomal insertions in *E. coli* cells were conducted by λRed recombination as described in (Bubnov et al. 2022). To obtain a double *soxR oxyR* mutation, the *cI-hok-strA* cassette was amplified from the pTmpSm plasmid with P6/P7 primers and integrated into the chromosome of *E. coli* BW25113 Δ*oxyR* cells. *E. coli* B233 strain with enhanced expression of *ftnA* was constructed by the insertion of a strong constitutive promoter P_H207_ (Deuschle et al. 1986) directly upstream of the *ftnA* open reading frame in the chromosome of *E. coli* MG1655. The insertional fragment P_H207out_-*cI-hok-neo* was amplified from the pTmpKmH207out plasmid with P8/P9 primers. To obtain a mutation in the *ryhB* regulatory RNA gene, the *cI-hok-neo* cassette was amplified from the pTmpKm plasmid with P14/P15 primers and integrated into the chromosome of *E. coli* BW25113 cells. *E. coli* MBM241-fecA and MBM243-ftnA strains were constructed by replacement of *cI-hok-neo* cassette in the chromosome of *E. coli* B2095 with the promoters of *fecA* or *ftnA* genes. Primers P16/P17 were applied to amplify P_*fecA*_ from *E. coli* MG1655gDNA and attach adapters. The fragment obtained was the template for amplification of the P_*fecA*_ insertional fragment using primers P18/P19. P_*ftnA*_ insertional fragment was amplified from pFtnA-lux plasmid with P18/P19 primers.

### 2.4. Chemicals

All chemicals were of analytical purity. Hydrogen peroxide (HP) was obtained from “Ferraine” (Russia). Methyl viologen (MV) was obtained from Sigma-Aldrich (St. Louis, MO, USA).

### 2.5. Biosensor luminescence measurement

Biosensor cells were grown in LB supplemented with appropriate antibiotics during 1-1.5 h until reaching an OD ∼ 0.1-0.2. Obtained cultures were divided into the 190 μL subcultures in separate wells of 96-well plates, then 10 μL samples of tested compound (HP, MV, or distilled water as negative control) were added. Cells were incubated without shaking at room temperature with regular measurements of total bioluminescence from each well during 2-3 h. Bioluminescence was measured using SynergyHT (Biotek Instruments, Winooski, VT, USA) with no filter, 135 gain and 2 s integration time. Luminescence values were expressed in relative light units (RLU), specific to the luminometer. Response amplitude (RA) for sample x was calculated using the formula:

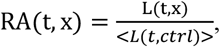

where L(t, x) – the luminescence of probe x in the time point t, <L(t,ctrl)> – the average luminescence at the time point t of negative control samples in at least 2 repeats (biosensor cells with addition of distilled water instead of HP or other supplements).

### 2.6. Measurement of intracellular iron content

For intracellular iron content analysis the ferrozine-based assay was used (Riemer et al. 2004). The volume of NaOH solution for bacteria lysis was adjusted for obtaining equal mass of dry cell per mL. The measurements of ferrozine-Fe^2+^ colored complex spectra were performed using Shimadzu UV-2450 (Kyoto, Japan) spectrophotometer. Iron concentration was then calculated from OD_562_ using the calibration with standard FeSO_4_ solutions.

## 3. Results

### 3.1. The P_ftnA_ promoter is activated by oxidative stress

To find out whether *ftnA* expression is activated in *E. coli* cells under oxidative stress, we used luminescent biosensor cells, the luminescence of which was determined by the expression of *luxCDABE* genes under the control of the *ftnA* gene promoter. To facilitate the study of the role of various genes in the regulation of *ftnA* expression, the pFtnA-lux biosensor plasmid with P_*ftnA*_-*luxCDABE* fusion was constructed. pFtnA-lux is derivative of pDEW201 promoter-probe vector (Van Dyk and Rosson 1998) obtained by insertion of *E. coli* P_*ftnA*_ promoter upstream of the *P. luminescens luxCDABE* genes. Upon activation of the P_*ftnA*_ promoter, *E. coli* cells harboring the pFtnA-lux plasmid produce more luminescence. The graphs in Figure 1 show the amplitude of the biosensor response over time to the addition of ferrous chloride (FeCl_2_), hydrogen peroxide (HP), and methyl viologen (MV). The response amplitude is calculated as a ratio of luminescence of the treated biosensor cells to the luminescence of untreated biosensor cells.

**Figure 1.**
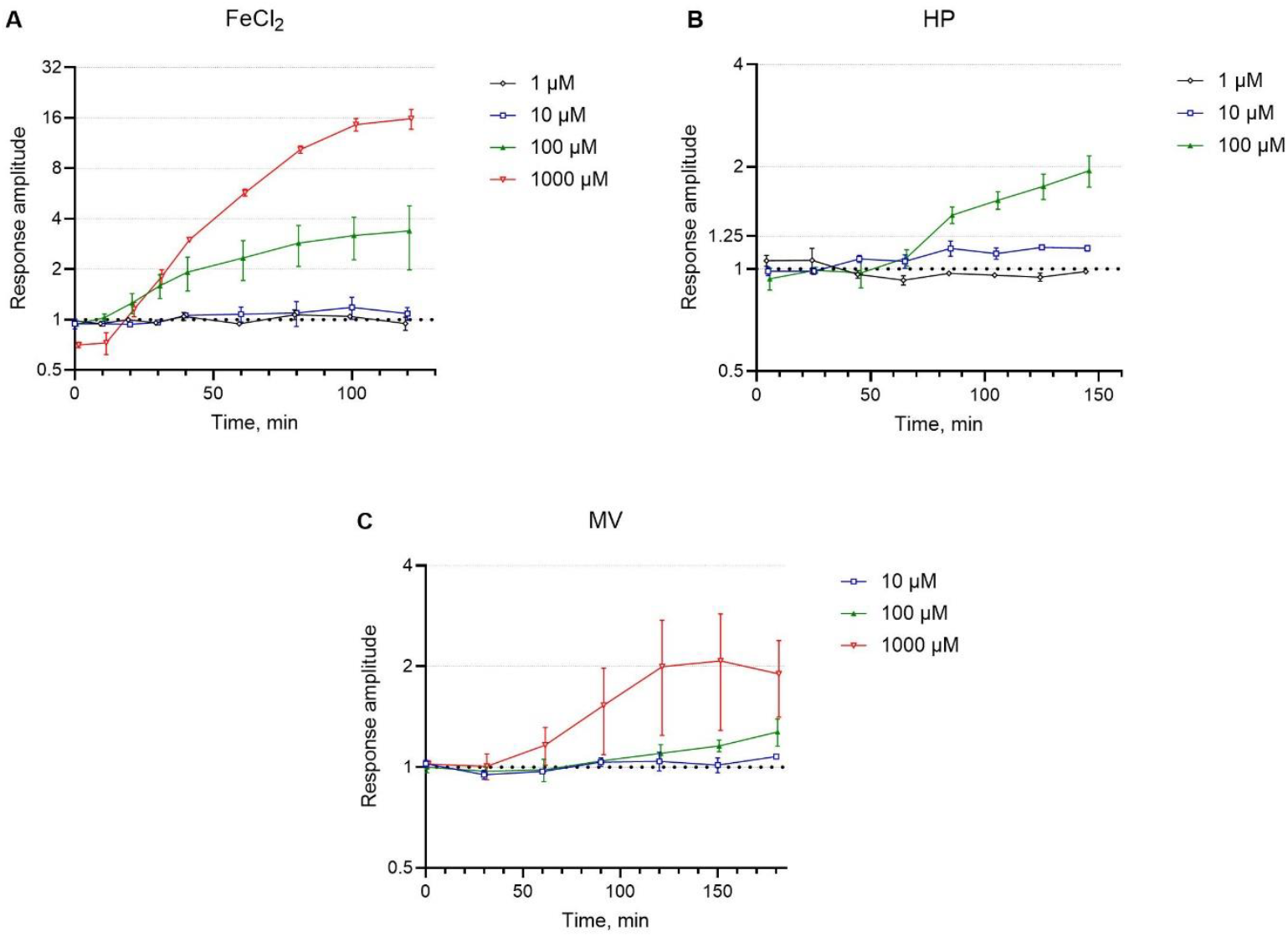
MG1655 pFtnA-lux biosensor reaction to the addition of Fe^2+^ in the form of FeCl_2_ (panels A), HP (panel B), or MV (panel C) to the culture medium. The graphs show the dependence of the response amplitude on time (ratio of the luminescence of a cell culture supplemented with FeCl_2_, HP, or MV to the luminescence of a cell culture without additions). Points and error bars represent mean and standard error of the mean calculated from 3 independent replicates. For the convenience of the reader, the curves are shifted relative to each other by 1 minute.

As one can see from the graphs shown in Figure 1, transcription from the P_*ftnA*_ promoter is activated when agents that cause oxidative stress are added. Significant enhancement in luminescence in response to the FeCl_2_ addition proves the functionality of the constructed biosensor, the main regulator of which is Fur, specific sensor of the Fe^2+^. The threshold concentration of added iron that can activate P_*ftnA*_ in LB-cultured cells is about 10 μM, which is close to the iron concentration in LB medium. Thus, we can judge that LB is not rich enough in iron for Fur to be maximally activated and cells are sensitive enough to detect fold changes in the concentration of iron.

When adding HP or MV, the maximum activation amplitude (from 1.5 to 3 folds) and the shortest response time (50-70 minutes) were approximately the same. However, the response of the biosensor to HP was much more stable, while with the addition of MV, both the time of promoter activation and the activation amplitude varied significantly from experiment to experiment, which is reflected in Figure 1C by large error bars. Thus, oxidative stress indeed leads to an activation of the P_*ftnA*_, but the work (Faulkner and Helmann 2011) described the effects emerging in only 50-70 minutes after oxidative stress exposure, when the peroxide should be neutralized.

### 3.2. P_ftnA_ activation after oxidative stress is determined solely by Fur and not by SoxR and OxyR

To study the mechanism of P_*ftnA*_ regulation under oxidative stress, we used *E. coli* strains with knockout mutations in the *fur, ryhB, soxR*, and *oxyR* genes, carrying the biosensor plasmid pFtnA-lux. Cells BW25113 Δ*fur* pFtnA-lux, BW25113 Δ*ryhB* pFtnA-lux, BW25113 Δ*oxyR* pFtnA-lux, BW25113 Δ*soxR* pFtnA-lux, BW25113 Δ*oxyR* Δ*soxR* pFtnA-lux were grown under the same conditions and exposed to HP at different concentrations. The obtained data on the kinetics of P_*ftnA*_ promoter activation in mutant strains are shown in Figure 2. The results for BW25113 pFtnA-lux do not differ from those for MG1655 pFtnA-lux given in Figure 1B.

**Figure 2.**
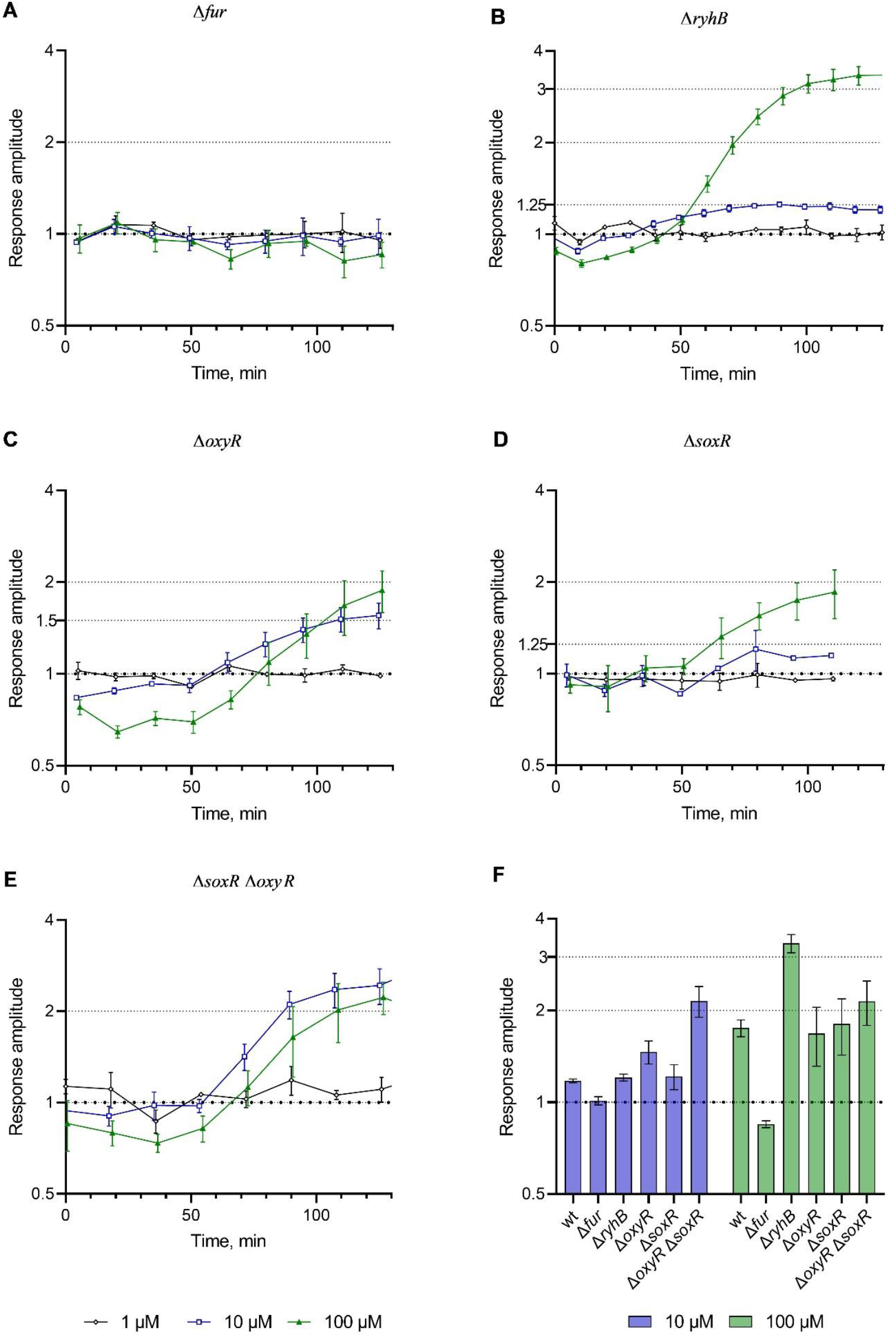
Response of the BW25113 pFtnA-lux cells with a mutation in *fur* (panel A), *ryhB* (panel B), *oxyR* (panel C), *soxR* (panel D), or both *soxR* and *oxyR* (panel E) regulatory genes to the addition of HP to the medium. Response amplitude of the biosensor cells at 120 min after HP treatment (panel F). “wt” corresponds to *E. coli* strain without listed mutations. Data presentation is equal to Figure 1.

As can be seen from the graphs in Figure 2, activation of P_*ftnA*_ after HP addition is determined by the Fur regulator, but not by OxyR or SoxR. The luminescence of BW25113 Δ*fur* pFtnA-lux cells in LB was initially ∼100 times lower than the luminescence of cells of the other listed strains and was not induced when HP was added (Figure 2A). Since P_*ftnA*_ is known to be regulated by competitive binding of the Fur-Fe^2+^ and H-NS proteins, it was rather expected. Less expected was the result for the cells with knockout of *ryhB* gene for regulatory RNA: a higher amplitude of P_*ftnA*_ activation was observed (Figure 2B).

Activation of P_*ftnA*_ by peroxide in *soxR* mutant cells (Figure 2D) is nearly the same to that in wild-type cells (Figure 1B). The mutation in the *oxyR* gene makes cells more vulnerable to HP, which is expressed in a sharp drop in the luminescence of biosensor cells shortly after the addition of 100 μM HP before the first measurement (Figures 2C and 2E). However, neither mutations in the individual *soxR* and *oxyR* genes, nor their combination lead to a decrease in HP-stimulated P_*ftnA*_ activation in *E. coli* cells. The double mutant becomes more sensitive and we can observe more pronounced activation of P_*ftnA*_ in response to HP at a concentration of 10 μM, which is near the lower limit of sensitivity of the biosensors based on OxyR- and SoxR-regulated promoters (Kotova et al. 2010).

### 3.3. Correlation between expression of ftnA and HP-mediated P_ftnA_ activation

The next question to answer was whether the activation of P_*ftnA*_ in response to the appearance of HP is a side effect of the activation of a number of intracellular regulatory cascades or FtnA is still involved in the cell reaction and the activation amplitude of its promoter reflects the cell’s needs for this protein. To do this, we used mutant *E. coli* strains: BW25113 Δ*ftnA* and B233 (P_H207_-*ftnA*), in which the chromosomal copy of the *ftnA* gene is either deleted (Baba et al. 2006) or placed under the control of the strong constitutive promoter P_H207_ (Deuschle et al. 1986). Increased expression of the *ftnA* gene in strain B233 was confirmed using RT-PCR (see Supplementary materials, Figures S1, S2). Cells of the listed strains were transformed with the pFtnA-lux plasmid. The transformants were treated with HP at various concentrations, and the amplitude of the biosensor response was measured (Figure 3).

**Figure 3.**
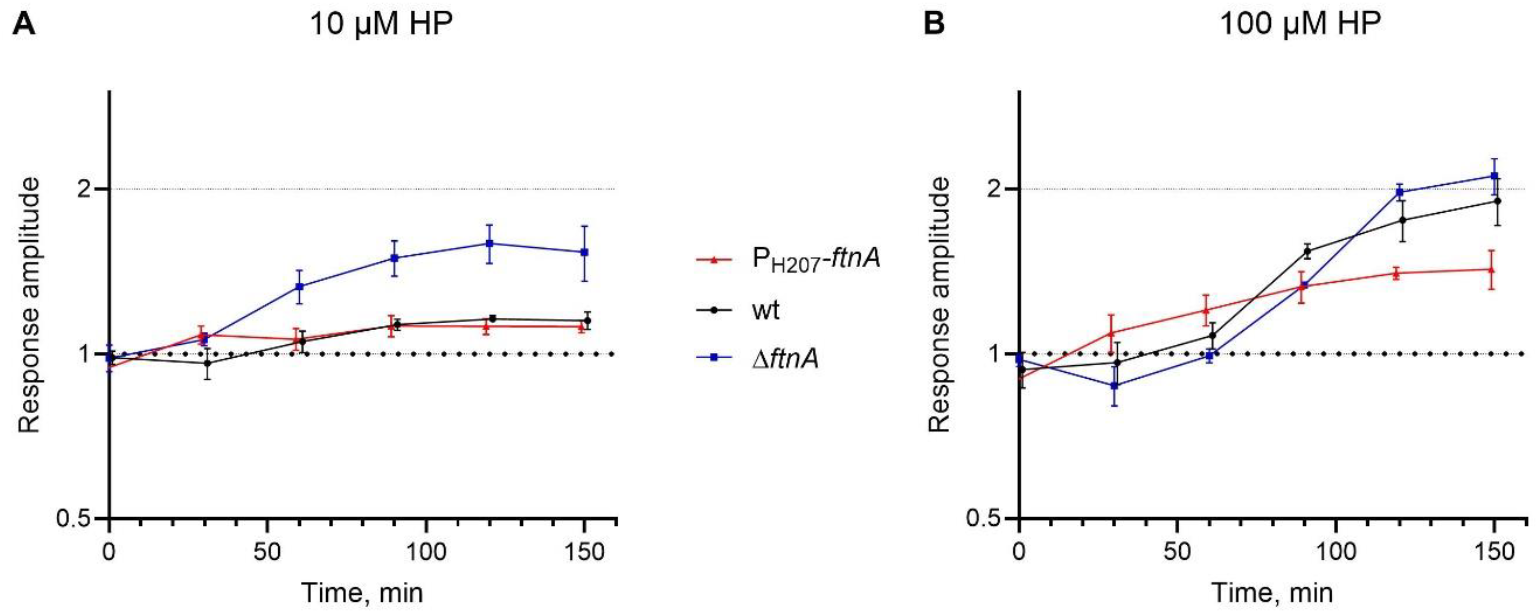
Response of the BW25113 Δ*ftnA* pFtnA-lux (“Δ*ftnA*”, blue squares), BW25113 pFtnA-lux (“wt”, black circles) and B233 pFtnA-lux (“P_H207_-*ftnA*”, red triangles) biosensor cells to the addition of HP to the medium at a concentration of 10 μM (panel A) or 100 μM (panel B).

As can be seen from the graphs in Figure 3, an increase in *ftnA* expression in the *E. coli* cell leads to a decrease in the amplitude of HP-induced activation of the P_*ftnA*_ promoter. The effect is more noticeable at an HP concentration of 10 μM. At 100 μM, two effects are observed from changes in the amount of FtnA in a cell. At the early stage, luminescence of the BW25113 Δ*ftnA* pFtnA-lux cells goes down, while luminescence of the other strains does not. Later, the amplitude of HP-induced activation of the P_*ftnA*_ promoter in Δ*ftnA* and wt strains exceeds that of the P_H207_-*ftnA* strain.

One possible explanation of these observations is that FtnA-producing cells (probable probiotic or biotechnological strains) are more sustainable to oxidative stress. To test this hypothesis, similar measurements were carried out using *E. coli* strains BW25113 Δ*ftnA* and B233 (P_H207_-*ftnA*), transformed with biosensor plasmids pSoxS, pDps, or pIscRSUA. These plasmids harbor the *P. luminescens luxCDABE* genes under the control of the P_*soxS*_, P_*dps*_, or P_*iscR*_ promoters. We didn’t observe any statistically significant dependence of the amplitude of HP-mediated activation of the oxidative stress promoters P_*soxS*_, P_*dps*_, and P_*iscR*_ on the level of *ftnA* expression (see Supplementary materials, Figure S3). It may indicate that FtnA is not directly involved in neutralizing oxidative stress. This conclusion is also supported by the results of investigation of the Δ*ftnA*, wt, and P_H207_-*ftnA* cells tolerance to HP-treatment. All three strains gave approximately the same results: 1 mM HP causes death of ∼70% cells in the culture, which was identified both by luminescence measurements right after HP addition and cell titer measurements (titration and plating were conducted in 30 min after HP-treatment); 100 μM HP did not cause significant changes in cell viability.

### 3.4. Oxidative stress leads to a sequential activation of iron uptake and iron storage genes

Since an increase in the amount of FtnA in a cell affected the regulation of P_*ftnA*_ under conditions of oxidative stress, but did not affect the activation of other promoters of the oxidative stress response system, experiments were conducted to determine the amount of iron in cells. Cells were grown to reach an OD of 0.05-0.1 in 40-100 mL LB, then divided into two equal portions and one of them was supplemented with HP. After 100 minutes of HP-treatment, cells were harvested from the suspension by centrifugation and the total content of intracellular iron was measured. The results of several independent measurements showed that specific iron content in cells (m/m) increases after HP treatment in comparison with the same cells in HP-free medium. This rise was 1-10% in the wild-type cells, and 30-60% in the strain that overproduces FtnA.

Taken together, all previous experiments lead us to the conclusion that HP-treatment causes changes in cellular iron concentration and Fur-mediated P_*ftnA*_ activation. Since our experiments use a multi-copy biosensor plasmid containing a DNA sequence for Fur binding, titration of this transcriptional repressor may contribute to the measured values of promoter activity change and affect the regulation of iron transport (Vassinova and Kozyrev 2000). To exclude the effect of titration, similar measurements of iron content were carried out using plasmid-free *E. coli* strains, and the same results of iron content rise were obtained. To verify the activation of P_*ftnA*_ in cells without Fur-box on multi-copy plasmid, the MBM243-ftnA strain was constructed. MBM243-ftnA is a derivative from MG1655 obtained by insertion of *P. luminescens luxCDABE* genes fused with *E. coli* P_*ftnA*_ promoter in *ara* locus in the chromosome.

To assess how oxidative stress affects iron uptake, we constructed a biosensor strain *E. coli* MBM241-fecA, which is analogous to MBM243-ftnA, but the expression of *luxCDABE* genes is controlled by the *fecA* gene promoter (Enz et al. 1995). Cells MBM241-fecA and MBM243-ftnA were grown to exponential phase and treated with various concentrations of HP or FeSO4, followed by luminescence measurements of the cell cultures (Figure 4).

**Figure 4.**
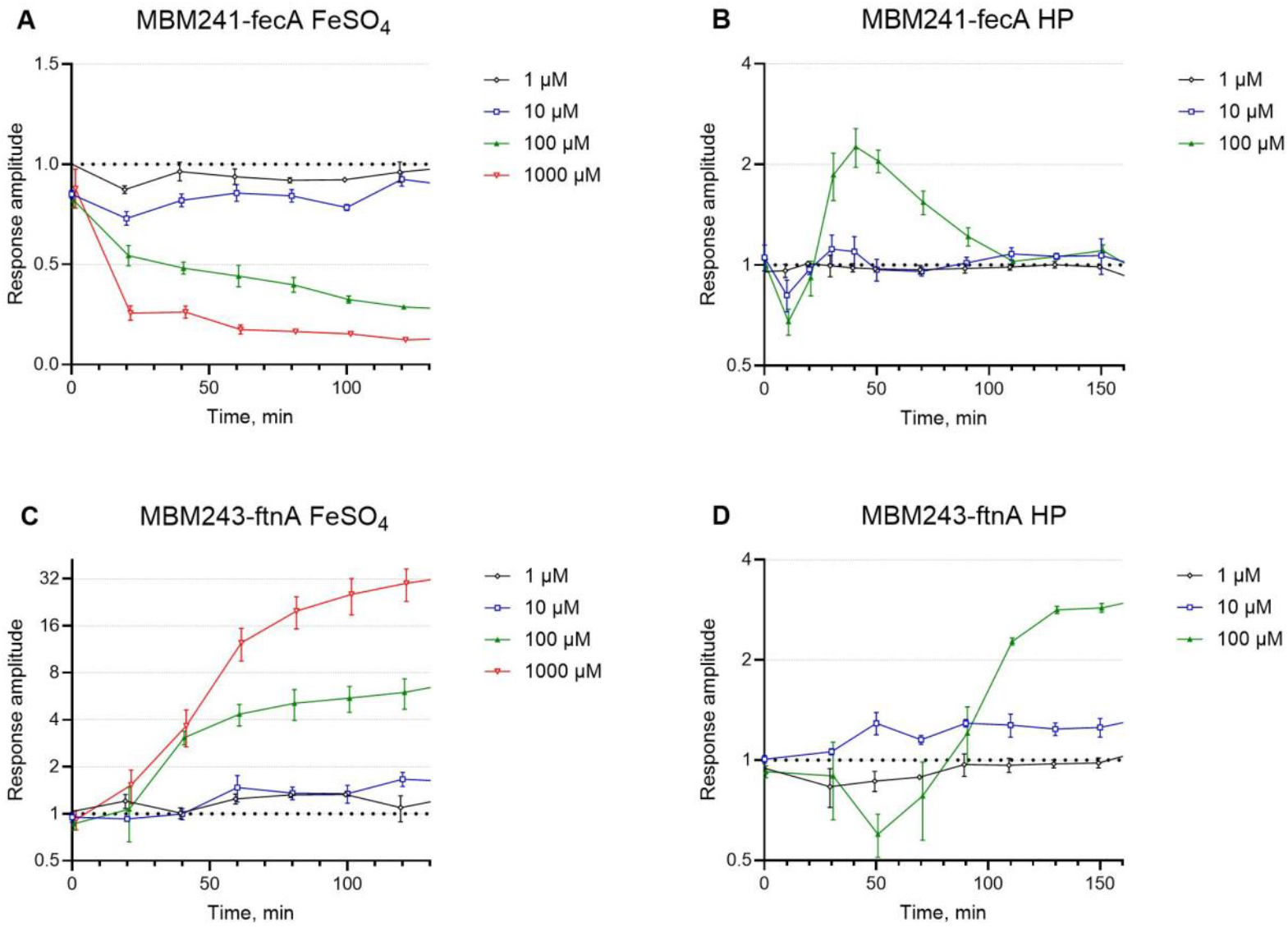
Response of the MBM241-fecA (panels A and B) and MBM243-ftnA (panels C and D) biosensor cells to FeSO_4_ (panels A and C) or HP (panels B and D) treatment.

Comparing the responses of plasmid-based biosensor (Figures 1A and 1B) and biosensor with chromosomally integrated promoter (Figure 4C and 4D), one can see that number of Furn-boxes or the P_*ftnA*_ promoter copies in a cell do not alter significantly iron- and HP-induced P_*ftnA*_ activation. The response amplitude of MBM243-ftnA biosensor cells is noticeably higher than that of MG1655 pFtnA-lux, but the other characteristics (threshold concentrations of HP, needed for activation, and time of response) are quite similar. Interestingly, lowered promoter copy-number and enhanced response amplitude allowed to observe repression of P_*ftnA*_ in 50 minutes in samples with 100 μM HP.

Graphs in the Figure 4 show P_*fecA*_ repression (panel A) and P_*ftnA*_ activation (panel C) shortly after FeSO_4_ treatment. This is in accordance with the logic that, in case of iron excess, cell needs to turn the iron uptake off and turn the iron sequestration on. Upon oxidative stress, the picture is different. Shortly after HP exposure, *fecA* promoter is induced (Figure 4B), indicating cell’s demand for Fe^2+^ and enhancement in expression of iron-uptake genes. Later, when oxidizing agent is neutralized, cellular Fe^2+^ concentration rises and transcription from P_*fecA*_ decreases. When adding 100 μM HP, in 20-50 minutes after HP-treatment one can see P_*fecA*_ activation simultaneously with P_*ftnA*_ repression (Figures 4B and 4D). It means that cells feel oxidative stress, Fur loses iron, and Fur-repressed promoters are activated, while the activity of the Fur-activated P_*ftnA*_ promoter is decreased. In 50-90 minutes after HP addition, cellular iron content rises, Fur binds Fe^2+^, and the P_*fecA*_ promoter is repressed, while P_*ftnA*_ is activated (Figures 4B and 4D).

## 4. Discussion

The main regulators of iron homeostasis in *E. coli* cells are Fur and RyhB. It was previously known that Fur, forming a complex with Fe^2+^, binds to the promoter region of *ftnA* and competitively replaces H-NS, thus removing repressor and enhancing transcription of *ftnA*. RyhB is a non-coding RNA regulated by the Fur repressor (Vassinova and Kozyrev 2000), which causes the rapid degradation of a number of mRNAs that encode iron-utilizing proteins. RyhB is not directly involved in P_*ftnA*_ regulation (Nandal et al. 2010). In (Massé and Gottesman 2002) Fur expression was shown to be positively regulated by the oxidative stress receptor systems OxyRS and SoxRS (Zheng et al. 1999). However, these data were not sufficient to determine how the expression of ferritin *ftnA* in *E. coli* cells would change under oxidative stress. On the one hand, the OxyR and SoxS proteins should enhance the transcription of *fur*, and with a sufficient amount of iron in the cell, an increase in the amount of Fur-Fe^2+^ and increased expression of *ftnA* can be expected (Bitoun et al. 2008; Faulkner and Helmann 2011). On the other hand, when peroxide appears in the cell, proteins inside the cell lose bound Fe^2+^, and free iron is oxidized to Fe^3+^, which should lead to the breakdown of the Fur-Fe^2+^ complex and its inactivation and, as a result, a decrease in *ftnA* expression (Blanchard et al. 2007; Cornelis et al. 2011; Pinochet-Barros and Helmann 2018).

LB medium contains ∼10 μM iron, however this amount is not enough to repress cellular iron uptake and activate iron storage. This conclusion can be drawn from the activation of P_*ftnA*_ in response to iron supplementation (Figures 1A and 4C) and the repression of P_*fecA*_ (Figure 4A). It means that in LB-cultured *E. coli* cells the amount of Fur-Fe^2+^ is not enough to occupy all Fur-boxes. On the other hand, the iron uptake could be activated, i.e. not all Fur-boxes are free. Thus, we can measure both activation and repression of Fur-regulated promoters in cells being grown in LB.

In this work, we showed that oxidative stress leads to an activation of the P_*ftnA*_ promoter in *E. coli* cells. From the experimental data, we are offering the model of FtnA-associated cellular processes driven by oxidative stress (Figure 5A). HP or another oxidizing agent activates OxyR and SoxR regulatory proteins and, on the contrary, inactivates Fur. Thus, promoters P_*soxS*_ and P_*dps*_, which are directly regulated by the oxidative stress sensors OxyR and SoxR, have large response amplitude (10-to 100-fold activation) and short time of response to HP and MV (5 to 15 minutes). At the same time, due to Fur inactivation, iron-uptake (P_*fecA*_) is activated and P_*ftnA*_ is repressed shortly after HP exposure. Release of iron from damaged cellular proteins and its simultaneous uptake from the medium lead to the rise of intracellular free iron. After HP neutralization, iron is being reduced and Fur is binding it, which leads to an activation of P_*ftnA*_ and a repression of promoters responsible for the iron-uptake (Figure 5B). This model explains the significant delay in the response of P_*ftnA*_ to HP (50-70 minutes) compared to the activation of OxyRS and SoxRS regulons (10-15 minutes).

**Figure 5.**
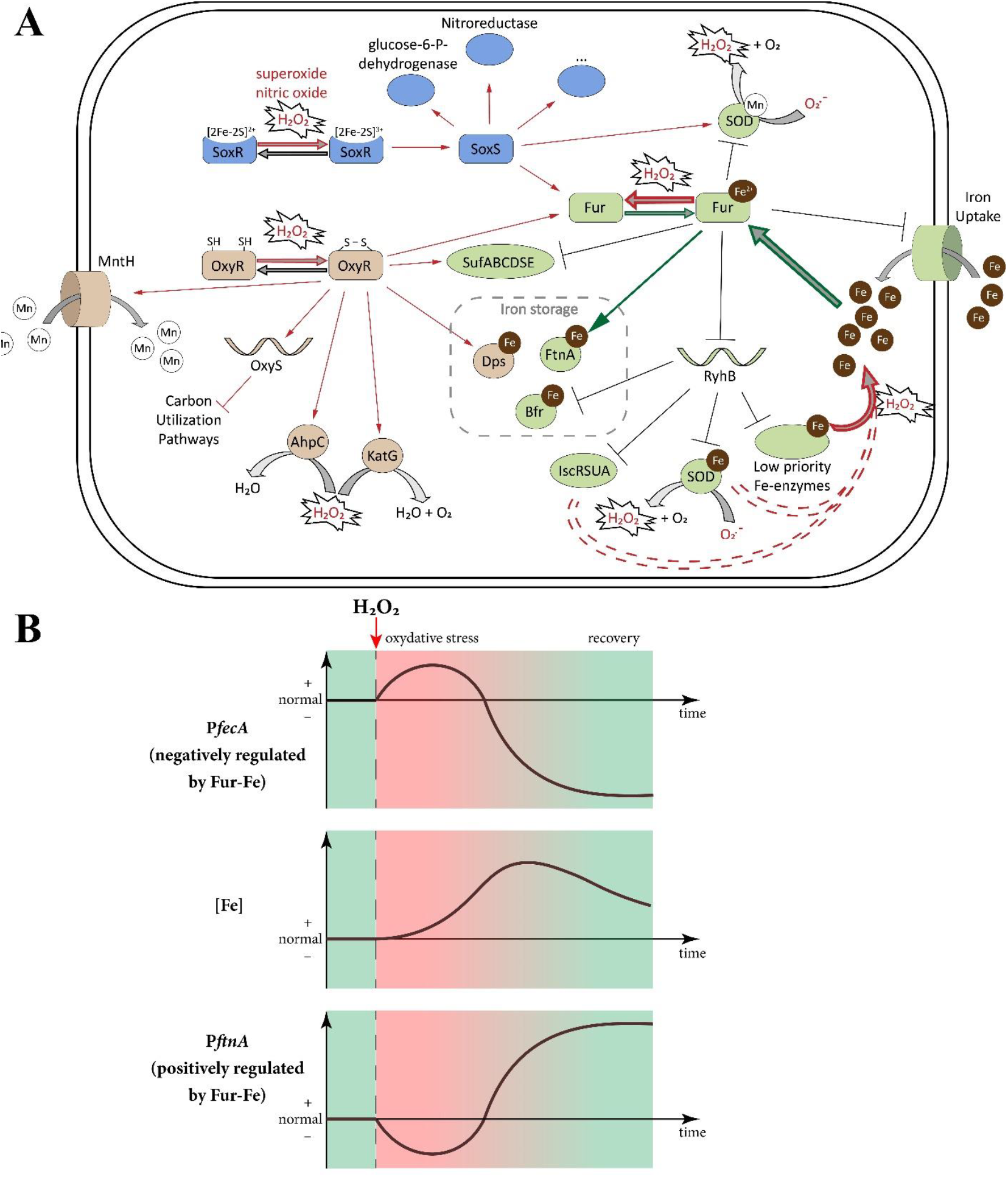
**A:** The OxyR, SoxRS, and Fur regulons in *E. coli*, revised version of scheme from (Faulkner and Helmann 2011). OxyR and SoxRS upregulates the synthesis of Fur under oxidative stress.

In Fe-rich conditions, Fur activates the expression of the FtnA, iron storage protein, by reversing H-NS-mediated silencing. However, Fur-Fe^2+^ is sensitive to H_2_O_2_ and can be inactivated upon exposure. Transcriptional and post-transcriptional regulation are shown by thin lines/arrows. Activation of ferritin expression is shown by a thicker arrow. Structural transformations, reactions, formation and disassembly of complexes, and ion transfers are shown by wide arrows with a gray center. Processes induced by oxidative stress are marked in red. Processes responsible for P_*ftnA*_ activation during the recovery after oxidative stress are marked in green. **B:** Scheme of sequential induction of P_*fecA*_ and P_*ftnA*_ after exposure to H_2_O_2_. Shortly after oxidative stress occurs, Fur-Fe is inactivated, *fecA* and other iron-uptake genes are activated, and iron concertation starts to grow. Later during cell recovery, iron concentration is high, Fur-Fe is assembled and active, *fecA* is being repressed, while *ftnA* – induced.

This model also explains why the amplitude of P_*ftnA*_ activation depends on the content of FtnA in the cell. Hypothesis of P_*ftnA*_ induction by the excess of free iron after HP exposure is supported by the fact that the response is amplified in an *ftnA* mutant and is diminished in an FtnA overproducing strain — indicating that iron sequestration blocks the response. SoxRS and OxyRS systems are not responsible for the P_*ftnA*_ activation upon oxidative stress neither directly (Figure 2A), nor via induction of *fur* expression (Figures 2C, 2D, and 2E). Probably the mechanism of OxyR- and SoxS-mediated *fur* induction (Zheng et al. 1999) is needed for better neutralization of iron-induced oxidative stress, but not for reaction on stress caused by other oxidizing agents. In the Δ*oxyR* Δ*soxR* double mutant, the amplitude of activation of P_*ftnA*_ by 10 μM peroxide is even higher than in the wild-type strain (Figure 2F). Since *dps* expression could not be induced by the oxidative stress in cells with Δ*oxyR* mutation, it can be assumed that Dps relieves iron fluctuations after the induction of oxidative stress by low concentrations of peroxide (about 10 μM). However, at higher concentrations of peroxide the *ftnA* induction was observed both in wild-type and mutant strains, so either Dps cannot sequester all exceeding iron, or active degradation of Dps by Clp proteases diminishes its role in an hour after HP exposure (Stephani et al. 2003). In addition, we observed an increase in the cellular iron content an hour after HP exposure, which is explained by the iron-uptake activation showed on the example of P_*fecA*_ (Figure 4B). Thus, results indicate the more prolonged effect of peroxide on the mutant cell (because of oxidative stress regulons malfunction), longer period of Fur and other proteins damaging leads to prolonged activation of iron uptake, and, as a result, higher iron concentration in a cell.

Works (Massé and Gottesman 2002; Massé et al. 2005) revealed some paradox with the regulation of *ftnA* and suggested that RyhB is involved in the regulation of *ftnA* not directly, but indirectly by activating the “iron sparing” mechanism. The “iron sparing” mechanism involves a rapid response of the cell to a deficiency of reduced iron in the composition of cellular proteins.

After inactivation of Fur, simultaneously with derepression of genes responsible for iron uptake, activation of RyhB synthesis leads to a decrease in iron consumption via mRNA degradation of many non-vital genes involved in iron metabolism (Jacques et al. 2006). This “iron sparing” mechanism should reduce fluctuations of iron concentration in a cell and ensure availability of iron for essential proteins. Therefore, we can see higher fluctuations in iron concentration in experiments with Δ*ryhB* mutants, since many iron-binding proteins are synthesized all the time. Under oxidative stress, these proteins are damaged, and therefore more iron is released in such cells than in cells with functional *ryhB* gene. While iron is actively pumped into the cell, these RyhB-regulated low priority enzymes consume Fe^2+^, it lowers effective Fe^2+^ concentration and makes cell to pump more iron. As a result, after some time P_*ftnA*_ is activated with larger amplitude in Δ*ryhB* mutants than in wild-type cells.

To summarize, oxidative stress leads to the postponed *ftnA* induction following the short-term repression. Moreover, this does not occur under the influence of the main regulators of the response to oxidative stress, but as a result of the changes in the free reduced iron content in the cell.

## Abbreviations

HP: hydrogen peroxide
MV: methyl viologen
RA: response amplitude
RLU: relative light units

## 5. Data availability statement

The data that supports the findings of this study is available from the corresponding author upon reasonable request.

## 6. Acknoledgements

Investigation of P_*ftnA*_ regulation in the *E. coli* cells under oxidative stress was supported by RSF 22-74-10047.

Assessment of influence of mutations on the sensitivity of P_*iscR*_-, P_*dps*_-, and P_*soxS*_-based biosensors was supported by the Ministry of Science and Higher Education of the RF, project FSMF-2023-0010.

The authors declare to have no conflict of interests.

## 7. Author contributions

V.O.M.: investigation, validation, data curation, visualization, writing – review & editing; A.D.G.: investigation, writing – review & editing; D.I.S.: investigation, writing – review & editing; F.V.V.: investigation, writing – review & editing; I.V.M.: conceptualization, supervision, data curation, formal analysis, writing – original draft, writing – review & editing, funding acquisition; S.V.B.: conceptualization, methodology, investigation, visualization, data curation, formal analysis, writing – original draft, writing – review & editing, funding acquisition.

## 8. Abbreviated Summary

Current models couldn’t predict the regulation of ferritin expression under oxidative stress. We showed for the first time that oxidative stress induced by different agents leads to a delayed induction of *ftnA* expression. It is independent of the function of oxidative stress response systems, SoxRS and OxyRS, but is entirely determined by the function of Fur. It could be explained by the significant changes in the level of iron in the cell after oxidative damage of the cell components.

