## Supplemental information 1 for "Oxidative stress leads to Fur-mediated activation of *ftnA* in *Escherichia coli* independently of OxyR/SoxRS regulators"

**Supplementary Material**

Supplementary Text: Verification of increased *ftnA* expression in strain B233 compared to MG1655.

Supplementary Figures S1-S3.

Supplementary Table S1.

**Text S1. Verification of increased *ftnA* expression in strain B233 compared to MG1655**

*E. coli* MG1655, B233 and BW25113 *ΔftnA* cells were grown in LB medium at 37 °C 200 rpm until OD_600_ ≈ 0.1 was achieved. When the specified optical density was reached, total RNA was isolated from the cells using the hot phenol-chloroform method (using Hot-Phenol) (Kumar et al. 2009). The concentration and quality of the isolated total RNA were analyzed using a NanoPhotometer ® P 330 spectrophotometer. Total RNA was additionally treated with DNase I (diaGene) during 10 min at 37°C with the following inactivation at 70°C during 10 min.

Reverse transcription polymerase chain reaction (RT-PCR) was performed on the isolated total RNA template using the OneTube RT-PCR SYBR kit (Eurogen, Russia). Primer pairs P10/P11 and P12/P13 were used to amplify regions of the *ftnA* (≈500 bp) and 16S rRNA (≈400 bp) genes, respectively (Table S1). Amplification was carried out in a Rotor-Gene Q amplifier, the data was processed by the Rotor-Gene 6000 program (Corbett Research, Australia). The amplification protocol was as follows: 50°C – 15 min; 95°C – 1 min; 40 cycles: 95°C – 15 s, 57°C – 20 s, 72°C – 30 s. Melting curves: 50°C – 1 min, from 50°C to 94°C in intervals of 5 s per 1°C.

RT-PCR amplification curves and gel electrophoresis of samples for the *ftnA* gene region after RT-PCR in 1% agarose gel are presented in Figures S1 and S2, respectively.

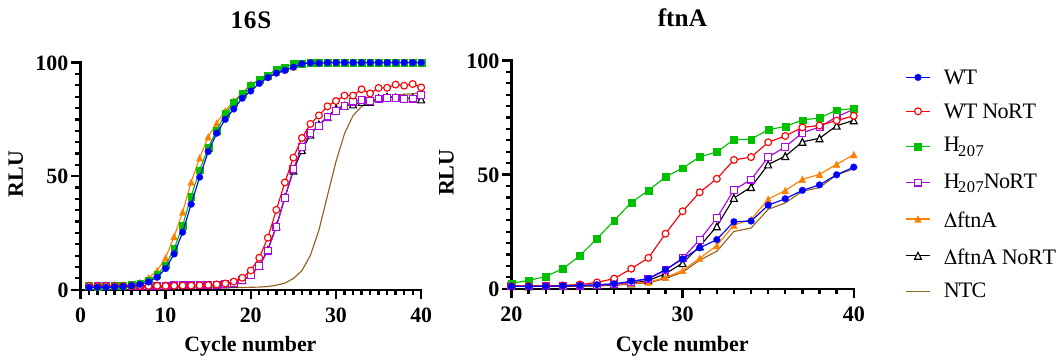

**Figure S1.** Amplification curves of 16S (left) and *ftnA* (right) gene fragments on template of total RNA, isolated from MG1655 (WT), B233 (P_H207_-*ftnA*), and BW25113 Δ*ftnA* (Δ*ftnA*) cells. Control samples designations: NoRT – without the addition of revertase, NTC – without the addition of a template (for each pair of primers).

| 100+bp Ladder (Evrogen) | WT | WT NoRT | H_207_ | H_207_ NoRT | Δ*ftnA* | Δ*ftnA* NoRT | NTC |
| --- | --- | --- | --- | --- | --- | --- | --- |

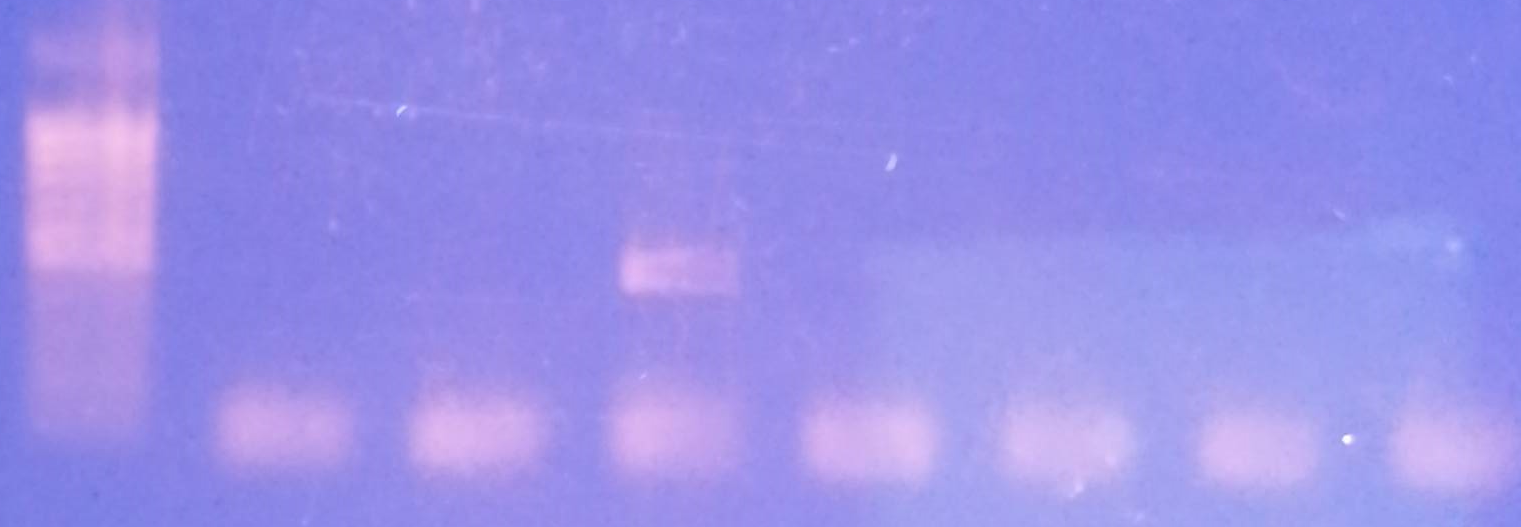

**Figure. S2.** Gel electrophoresis of RT‑PCR products, obtained with primers for amplification of *ftnA*.

As can be seen from Figures S1 and S2, transcription of the *ftnA* gene under the control of the P_H207_ promoter is significantly higher than in the wild-type strain MG1655. The absence of a specific product in the sample obtained using RNA from BW25113 ∆*ftnA* cells is explained by the absence of the corresponding gene in the cell. The absence of a specific product in the sample obtained using RNA from MG1655 cells can be explained by the very low level of *ftnA* expression, which is in agreement with previously published work (Massé and Gottesman 2002).

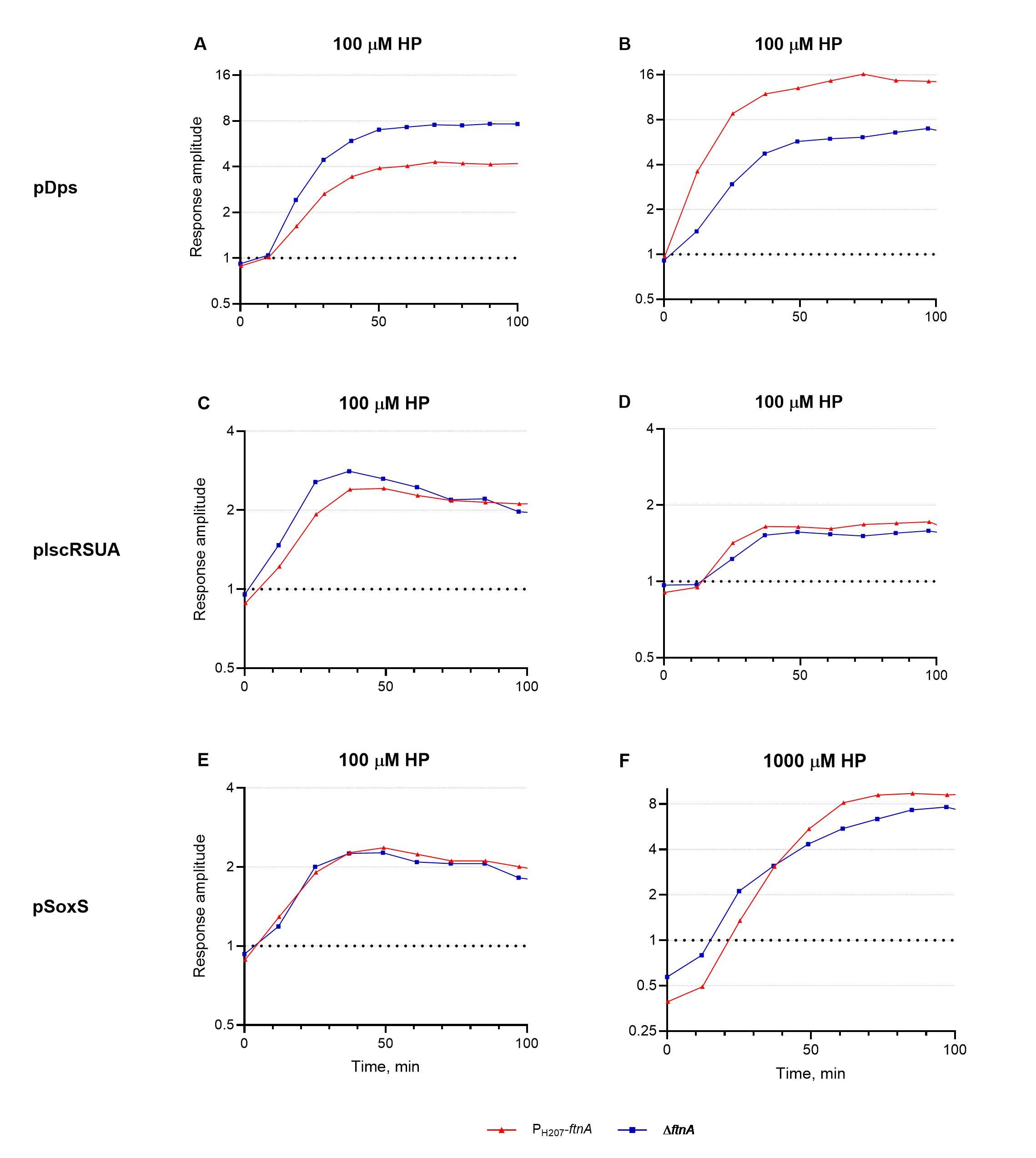

**Figure S3.** Response of the BW25113 ∆*ftnA* (“∆*ftnA*”, blue squares) and B233 (“P_H207_-*ftnA*”, red triangles), transformed with biosensor plasmids pDps (panels A, B), pIscRSUA (panels C, D), or pSoxS (panels E, F), to the addition of HP to the medium. The graphs show the dependence of the response amplitude on time (ratio of luminescence of a cell culture supplemented with HP to luminescence of a cell culture without it).

**Table S1.** Primers used in the study.

| **Name** | **Sequence 5’ → 3’** | **Note** |
| --- | --- | --- |
| P1 | TGCCAGGAATTGGGGATCGGTTCACTTTGTCATATCAGTGCCATCA | For plasmid pFtnA-lux construction |
| P2 | CGCGAGCTCGGTACCCGGGTGAAGAGTACAGTTCCAGGTTCATC |  |
| P3 | GTCTCATGTCTTACTTCACCTCAA | For plasmid pIscRSUA construction |
| P4 | GGTAACAAGGCCGAATAACAG |  |
| P5 | TCGGGAAAGATTTCAACCTGGCCGTTAATA | For sequencing of insertions in the pDEW201 |
| P6 | AGTTAACTTGAGGTAAAGCGATTTATGGAAAAGAAATTACCCCGCATTAATTAATTTTTCGAAAAAAGGCCTCCCAA | For amplification of *cI-hok-strA* cassette for knockout of *soxR* |
| P7 | GACAAAGACCGGAAAACAAACTAAAGCGCCCTTGTGGCGCTTTAGTTTTGATGCCATCTGGTATCACTTA |  |
| P8 | ATTTCTGGTTTCAGCATGATAGTGCTCCACAAAGGTTATATTTGTTGCTGTGGCGTCTTTTTGTTTATCAATTTATGAAGTATAaTATACGAGAATTTGAAGCGTTTAGCAAATG | For amplification of *P_H207out_-cI-hok-neo* cassette for transcriptional fusion of P_H207_ with *ftnA* |
| P9 | CATATCAGTGCCATCATTCCGCCTTCCCTGTCTTTCTCTAAATATTTTAATCAGCGACCATCTGGTATCACTTAAAGGTATTA |  |
| P10 | TTAGTTTTGTGTGTCGAGGGTAGAGAGTTC | RT-PCR on *ftnA* |
| P11 | GAAATGATTGAAAAACTTAATGAGCAGATGAAC |  |
| P12 | CGTGCCAGCAGCCGCGGTAATA | RT-PCR on 16S rRNA |
| P13 | GGCCCCCGTCAATTCATTTGAGT |  |
| P14 | CGCATAACGAACACAAGCACTCCCGTGGATAAATTGAGAACGAAAGATCATTTTTCGAAAAAAGGCCTCCCAAA | For amplification of *cI-hok-kanR* cassette for *ryhB* knockout |
| P15 | TTTGCAAAAAGTGTTGGACAAGTGCGAATGAGAATGATTATTATTGTCTCATGCCATCTGGTATCACTTAAAGG |  |
| P16 | CTGCCAGGAATTGGGGATCGGTTCAGTCTATTACGCGGTACT | For P*_fecA_* amplification |
| P17 | GGCGCGAGCTCGGTACCCGGGAGGCTCAGCGAATGGTGTTAACCAAAG |  |
| P18 | ATCAGCCAAACGTCTCTTCAGGCCACTGACTAGCGATAACTTTCCCCACATAAACTGCCAGGAATTGGGGATC | For transferring of promoters from pDEW201-based plasmids into *E. coli* B2095 chromosome |
| P19 | TAGTCATATTTGCCCCAACAGTTGCGCAGCCTGAATGGCGCGAGCTCGGTACCCGG |  |
